# From toxic waste to beneficial nutrient: acetate boosts *E. coli* growth at low glycolytic flux

**DOI:** 10.1101/2022.09.20.506926

**Authors:** Pierre Millard, Thomas Gosselin-Monplaisir, Sandrine Uttenweiler-Joseph, Brice Enjalbert

## Abstract

Acetate, a major by-product of glycolytic metabolism in *Escherichia coli* and many other microorganisms, has long been considered a toxic waste compound that inhibits microbial growth. This counterproductive auto-inhibition represents a major problem in biotechnology and has puzzled the scientific community for decades. Recent studies have revealed that acetate is also a co-substrate of glycolytic nutrients and a global regulator of *E. coli* metabolism and physiology. Here, we used a systems biology strategy to investigate the mutual regulation of glycolytic and acetate metabolism. Computational and experimental results demonstrate that reducing the glycolytic flux enhances co-utilization of acetate with glucose. Acetate metabolism thus compensates for the reduction in glycolytic flux and eventually buffers carbon uptake so that acetate, far from being toxic, actually enhances *E. coli* growth under these conditions. We validated this mechanism using three orthogonal strategies: chemical inhibition of glucose uptake, glycolytic mutant strains, and alternative substrates with a natively low glycolytic flux. Acetate makes *E. coli* more robust to glycolytic perturbations and is a valuable nutrient, with a beneficial effect on microbial growth.

## Introduction

Metabolism is the fundamental biochemical process that converts nutrients into energy and cellular building blocks. The preferred nutrients of many organisms are glycolytic substrates, such as glucose, which are oxidized into pyruvate by glycolysis. In the presence of oxygen, pyruvate can then be fully oxidized into carbon dioxide and water via respiratory pathways involving the tricarboxylic acid (TCA) cycle and oxidative phosphorylation. When oxygen is limiting, pyruvate is incompletely oxidized through non-respiratory pathways, and fermentation by-products are released into the environment. Despite fermentation being less efficient than respiration, fermentation by-products are still released in the presence of oxygen by most microorganisms, including bacteria (Harden, 1901), yeasts (Pasteur, 1857) and mammalian cells (Warburg *et al*, 1927).

Acetate is one of the most common by-products of glycolytic metabolism in *Escherichia coli* and many other microorganisms (Harden, 1901; Kutscha & Pflügl, 2020; Trcek *et al*, 2015). The established view is that acetate and other fermentation by-products are toxic and inhibit microbial growth (Beck *et al*, 2022; Enjalbert *et al*, 2017; Hollowell & Wolin, 1965; Kutscha & Pflügl, 2020; Luli & Strohl, 1990; Pinhal *et al*, 2019; Roe *et al*, 1998; Roe *et al*, 2002; Russel, 1992; Trcek *et al*., 2015; Warnecke & Gill, 2005). Why *E. coli* should produce a self-toxic by-product is an intriguing question, particularly since *E. coli*’s natural environment, the gut, is acetate-rich. Several explanations for the toxic effect of acetate have been proposed, the classical one being the uncoupling effect of organic acids (Russel, 1992), with acetic acid freely diffusing through the membrane before dissociating intracellularly into acetate and a proton due to its low pKa, and the energy required to expel the excess protons from the cell to maintain pH homeostasis being detrimental to growth. Another hypothesis is that the presence of acetate in the cytoplasm may affect the anion balance of the cell (Roe *et al*., 1998), reducing the pool of other anions – primarily glutamate – available to maintain osmotic pressure and electroneutrality and thereby impeding cell function. Acetate has also been reported to inhibit a step in the methionine biosynthesis pathway (Roe *et al*., 2002), reducing the intracellular methionine pool and concomitantly increasing the concentration of the toxic intermediate homocysteine. Finally, acetate may modulate the accumulation of acetyl-phosphate (Weinert *et al*, 2013; Wolfe *et al*, 2003), an intermediate of the acetate pathway and a signaling metabolite that can phosphorylate and acetylate enzymes and regulatory proteins with broad physiological consequences. However, none of these studies conclusively explains how acetate inhibits microbial growth (Pinhal *et al*., 2019). The physiological role of fermentation under aerobic conditions also remains unclear, especially since it leads to the accumulation of toxic by-products at the expense of biomass production.

Acetate overflow and its impact on *E. coli* are also important issues in biotechnology, where acetate is both produced from glycolytic carbon sources and found in many renewable resources (Klinke *et al*, 2004; Kutscha & Pflügl, 2020; Palmqvist & Hahn-Hägerdal, 2000; Sun *et al*, 2021; Trcek *et al*., 2015). As well as inhibiting growth, acetate production diverts carbon that could otherwise be used to synthesize biomass or valuable compounds. Acetate therefore decreases the productivity of bioprocesses. Since acetate is only produced at high glucose uptake rates (Nanchen *et al*, 2006; Renilla *et al*, 2012; Valgepea *et al*, 2010), a common approach to reduce acetate overflow is to limit glucose uptake by process engineering (e.g. glucose-limited chemostat or fed-batch culture) or by metabolic engineering (e.g. deletion of glucose transporters) (Bernal *et al*, 2016; De Mey *et al*, 2007; Eiteman & Altman, 2006; Fuentes *et al*, 2013; Kutscha & Pflügl, 2020; Luli & Strohl, 1990; Peebo *et al*, 2014). The utilization of acetate in combination with glycolytic substrates could alleviate growth inhibition and allow acetate to act as both a renewable carbon source and a relevant (co-)substrate for bioproducts that are derived from acetyl-CoA (Krivoruchko *et al*, 2015; Kutscha & Pflügl, 2020; Peebo *et al*., 2014; Wei *et al*, 2013). However, despite decades of attempts to reduce its accumulation and improve its utilization, acetate is still considered a major problem in biotechnology (Bernal *et al*., 2016; De Mey *et al*., 2007; Luli & Strohl, 1990; Pinhal *et al*., 2019).

Recently, we found that acetate can also act as a co-substrate for glycolytic carbon sources such as glucose, fucose and gluconate (Enjalbert *et al*., 2017), and is therefore more than just a waste by-product. The switch between acetate production and consumption was found to be determined thermodynamically by the acetate concentration itself. Acetate is also a global regulator of *E. coli* metabolism and physiology (Millard *et al*, 2021). These observations suggest an alternative rationale for acetate toxicity, namely that the inhibition of growth and glucose uptake may result from coordinated downregulation of the expression of most central metabolic genes (glucose uptake systems, glycolysis, TCA cycle) and ribosomal genes. To date, the role of acetate as a co-substrate and regulator of glycolytic metabolism has only been investigated under high glycolytic flux, and little is known about the response of *E. coli* to acetate at low glycolytic fluxes, conditions that are nevertheless commonly experienced by this bacterium in its natural, industrial and laboratory environments (Cummings & Englyst, 1987; Cummings *et al*, 2001; de Graaf *et al*, 2010; Fabich *et al*, 2008; Jones *et al*, 2007; Kleman & Strohl, 1994; Macfarlane *et al*, 1992; Shen *et al*, 2011; Wolfe, 2005).

In this study, we took a systems biology perspective to clarify the interplay between glycolytic and acetate metabolism in *E. coli*. We used a kinetic model of *E. coli* metabolism to generate hypotheses on the relationships between glycolytic, acetate and growth fluxes. We validated model predictions experimentally, explored the underlying biochemical and regulatory mechanisms on glucose, and extended our findings to other glycolytic nutrients.

## Results

### Model-driven hypotheses on the interplay between glucose and acetate metabolism in *Escherichia coli*

We explored the functional relationship between glycolytic and acetate metabolism in *Escherichia coli* using a coarse-grained kinetic model coupling glucose and acetate metabolism with growth (Figure 1A) (Millard *et al*., 2021). The model included two compartments, six compounds, and six reactions representing (i) glucose uptake and its conversion into acetyl-CoA by glycolysis, (ii) acetyl-CoA utilization in the TCA cycle and anabolic growth pathways, and (iii) the reversible conversion of acetyl-CoA into acetate via phosphotransacetylase (Pta), acetate kinase (AckA), and acetate exchange between the cell and its environment. Acetate utilization via acetylCoA synthetase (Acs), which is not active when glucose is present (Enjalbert *et al*., 2017), was not included. The model also included two inhibitory terms representing on the one hand inhibition of glucose uptake and glycolysis, and on the other inhibition of the TCA cycle and growth, in line with the observed inhibition of catabolic and anabolic processes (Millard *et al*., 2021). This model has been extensively validated and shown to have good predictive capabilities (Millard *et al*., 2021). We used it to simulate the response of glycolytic, acetate and growth fluxes to different degrees of glycolytic limitation by progressively reducing the activity (i.e. Vmax) of the glycolytic pathway – and thereby the glycolytic flux – from 100 % to 20 % of its initial value. Since glycolytic and acetate fluxes are both determined by the concentration of acetate (Enjalbert *et al*., 2017; Millard *et al*., 2021; Pinhal *et al*., 2019), these simulations were carried out over a broad range of acetate concentrations (from 0.1 to 100 mM).

**Figure 1.**
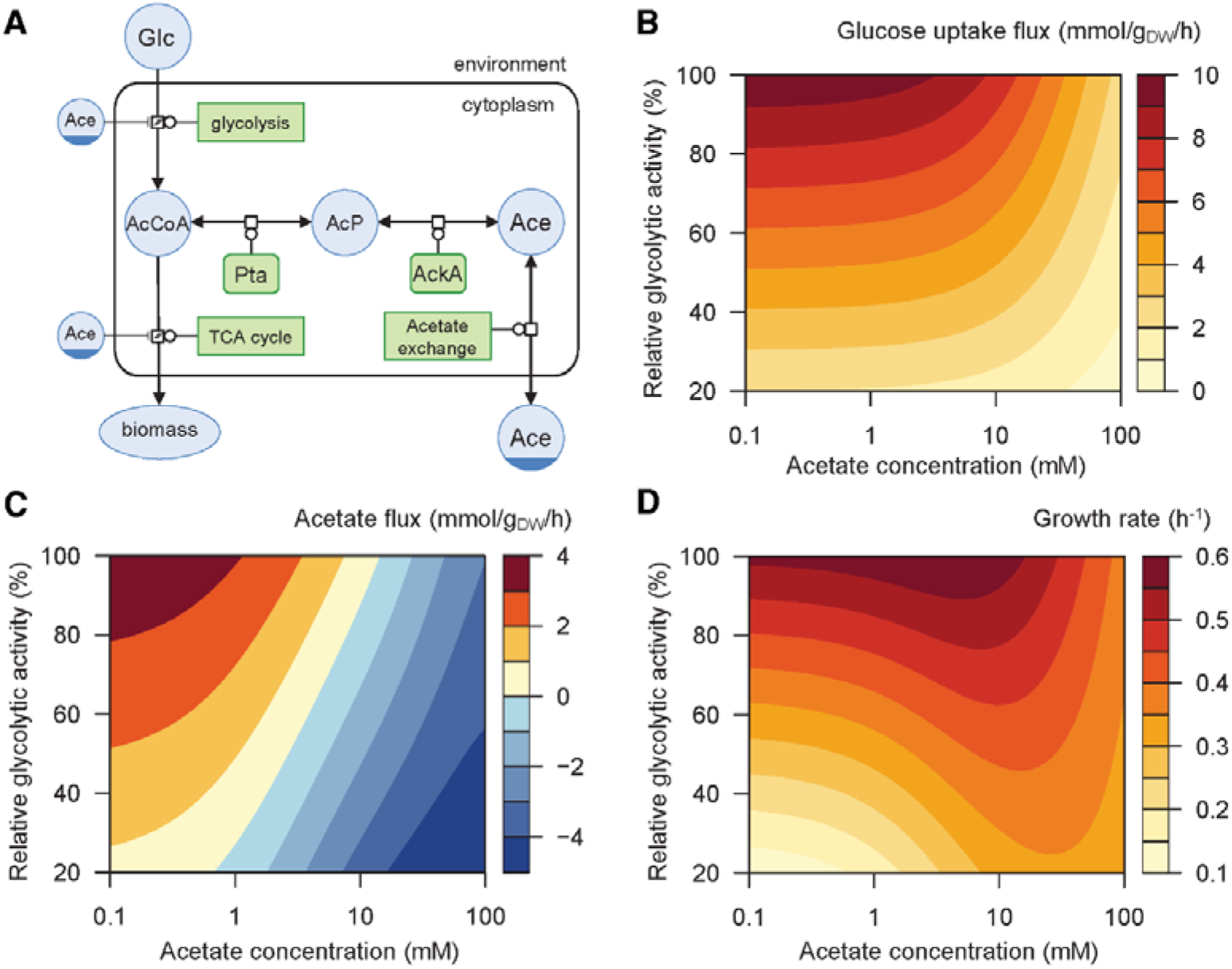
Predicted response of *E. coli* to glycolytic and acetate perturbations. A Representation (in Systems Biology Graphical Notation format, http://sbgn.org) of the glucose and acetate metabolism of *Escherichia coli*, as implemented in the kinetic model used in the study (reproduced from (Millard *et al*., 2021)). B-D Glucose uptake flux (B), acetate flux (C) and growth rate (D) simulated over a broad range of glycolytic activity levels (from 20 to 100 % of the initial Vmax) and acetate concentrations (from 0.1 to 100 mM).

The results obtained (Figure 1B-D) highlight the strong bidirectional interplay between glycolytic and acetate metabolism, which eventually modulate growth. Glucose uptake and acetate flux both respond nonlinearly to perturbations (Figure 1B-C). According to the model, increasing the acetate concentration inhibits glycolysis (Figure 1B) and reverses the acetate flux (Figure 1C) over the full range of glycolytic fluxes, as previously reported at the highest glucose uptake rate (Millard *et al*., 2021). The acetate concentration at which the flux reversal occurs appears to depend on the glycolytic flux (Figure 1C). Reducing the glycolytic flux lowers acetate production when acetate is a by-product of glucose metabolism, but seems to enhance acetate utilization when acetate is a co-substrate of glucose (Figure 1C). Finally, the glycolytic flux and the acetate concentration both have a strong, nonlinear effect on the growth rate (Figure 1D). At maximal glycolytic activity, the predictions of the model are consistent with the established deleterious effect of acetate on microbial growth (Figure 1D). However, despite the dual inhibition of catabolism and anabolism by acetate included in the model, it also predicts that at low glycolytic flux, increasing the acetate concentration should boost *E. coli* growth (Figure 1D). A global sensitivity analysis showed that the predictions of the model are robust to parameter uncertainty. We present these model-driven hypotheses in detail (including the sensitivity analysis results) in the following sections along with experimental results collected for validation. Note that none of the data obtained in this study were used to build or calibrate the model and therefore represent independent tests of its predictions and inferred hypotheses.

### Enhanced acetate utilization at low glycolytic flux buffers carbon uptake

We investigated the glycolytic control of *E. coli* metabolism by confronting the model predictions with experimental data (Figures 1 and 2). The simulations show that, at low acetate concentrations (0 mM in Figure 2A-D), acetate production is predicted to increase with the glycolytic flux, in line with the relationship observed in glucose-limited chemostat experiments (Nanchen *et al*., 2006; Renilla *et al*., 2012; Valgepea *et al*., 2010) and in a set of mutants with different levels of glucose uptake (Fuentes *et al*., 2013). In contrast, when the acetate concentration is high (30 mM in Figure 2A-D), such that it is co-consumed with glucose, acetate utilization is expected to increase when the glycolytic flux decreases. This response implies, as suggested previously (Millard *et al*., 2021) but never verified experimentally, that the control exerted by glycolysis on the acetate flux is non-monotonic.

**Figure 2.**
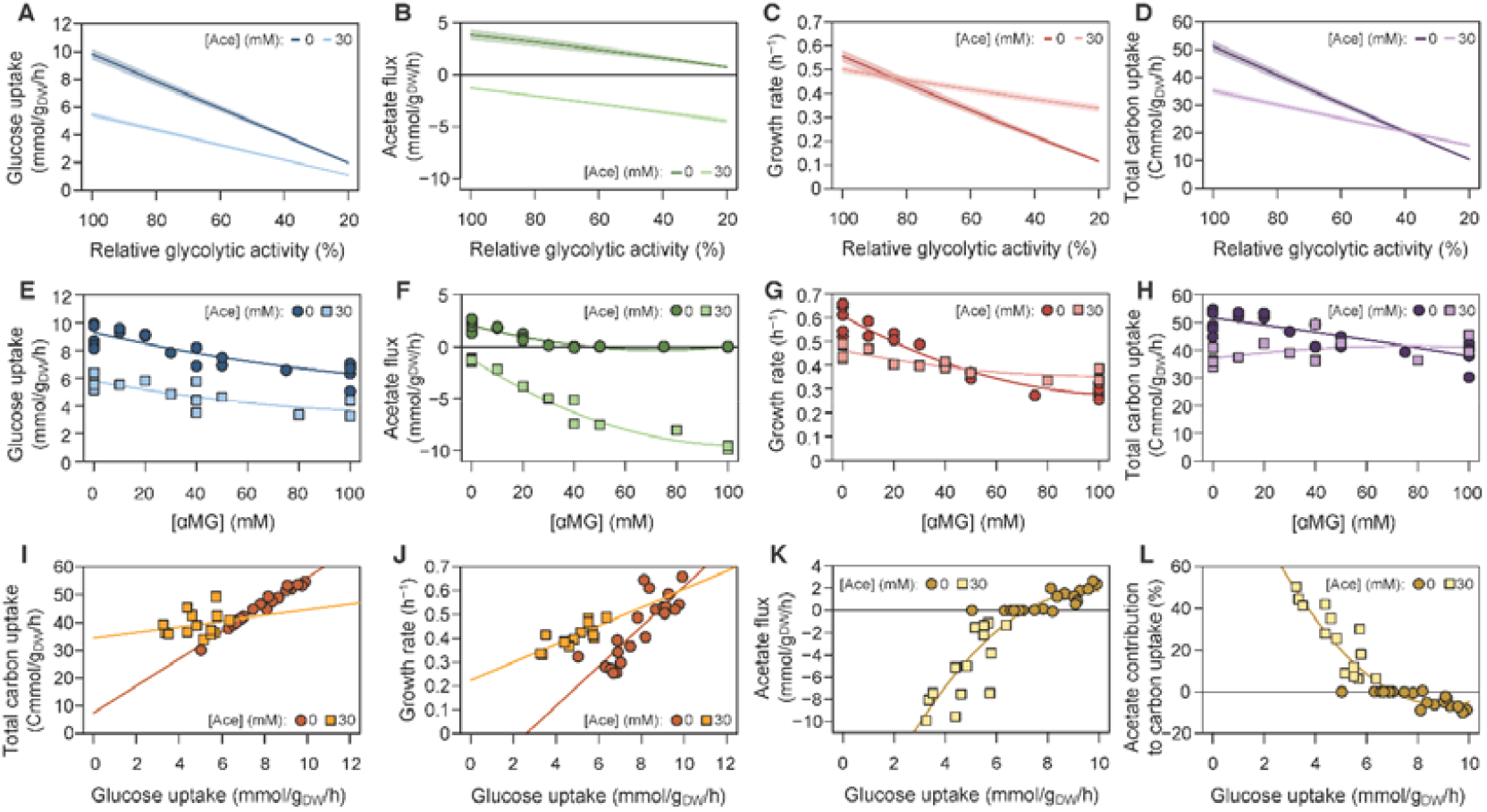
Glycolytic control of *E. coli* metabolism. A-D Predicted glucose uptake flux (A), acetate flux (B), growth rate (C), and total carbon uptake flux (D) of *E. coli* on glucose (15 mM) as a function of the glycolytic activity (20–100 % of the maximal activity), at low (0 mM) and high (30 mM) acetate concentrations. E-H Experimental glucose uptake flux (E), acetate flux (F), and growth rate (G) of *E. coli* grown on glucose (15 mM) with different concentrations of αMG (0–100 mM), at low (0 mM) and high (30 mM) acetate concentrations. These data were used to quantify the total carbon uptake flux as a function of the αMG concentration (H). Each data point represents an independent biological replicate, and lines represent the best polynomial fits. I, J Total carbon uptake (I) and growth rate (J) as a function of the glucose uptake flux, at low (0 mM) and high (30 mM) acetate concentrations. Each data point represents an independent biological replicate, and lines represent the best linear fits. K, L Relationships between the acetate flux and glucose uptake (K), and between the contribution of acetate to carbon uptake and glucose uptake (L). Each data point represents an independent biological replicate, and lines represent the best polynomial fits.

To test this hypothesis, we measured growth, glucose uptake, and acetate fluxes in *E. coli* K-12 MG1655 grown on 15 mM glucose and increased the concentration of the glycolytic inhibitor methyl α-D-glucopyranoside (αMG, an analogue of glucose that is taken up and phosphorylated but not metabolized by glycolysis) (Bren *et al*, 2016; Negrete *et al*, 2010) (Figure 2E-H). Experiments were carried out without acetate or with 30 mM acetate for comparison with the predictions shown in Figure 2A-D. Results confirm that αMG inhibits glucose uptake under both conditions (Figure 2E). In the absence of acetate, the acetate (production) flux decreased when the αMG concentration was increased (Figure 2F), as expected based on the positive control exerted by glycolysis on the acetate flux. In contrast, in the presence of 30 mM acetate, increasing the αMG concentration increased the acetate uptake flux (Figure 2F). These results are consistent with the predictions of the model (Figure 2A-D) and demonstrate that glycolysis controls the acetate flux in a non-monotonic manner. The control exerted by glycolysis on acetate metabolism thus depends on the concentration of acetate itself.

The growth and total carbon uptake rates decreased when the αMG concentration was increased in the absence of acetate (Figure 2G-H), but were more stable in the presence of acetate despite a ∼2-fold reduction in glucose uptake (between 5.7 and 3.3 mmol·g_DW_^−1^·h^−1^ with 30 mM acetate; Figure 2E). Indeed, both the total carbon uptake and the growth rates were less sensitive to glycolytic perturbations in the presence of acetate than in the absence of acetate (Figure 2I-J), suggesting that acetate compensates for the drop in glucose uptake. Consistently, a strong, nonlinear relationship between glucose uptake and acetate flux was observed in all experiments (Figure 2K), which was reflected in the contribution of acetate to the total carbon uptake of *E. coli* (Figure 2L). At the maximal glucose uptake rate, acetate production represented a 10 % loss in carbon. However, when glucose uptake was below 7 mmol·g_DW_ ^−1^·h^−1^, acetate was co-utilized with glucose and contributed positively to the carbon balance (up to 50 % at a glucose uptake rate of 3.3 mmol·g_DW_^−1^·h^−1^; Figure 2L). These results highlight the coupling between glycolytic and acetate fluxes in *E. coli*, with acetate buffering carbon uptake at low glycolytic flux, thereby improving the robustness of *E. coli* to glycolytic perturbations.

### The glycolytic flux determines the acetate concentration threshold at which the acetate flux reverses

As already reported at maximal glucose uptake rates (Enjalbert *et al*., 2017; Millard *et al*., 2021), the acetate concentration threshold at which the acetate flux switches between production and consumption is about 10 mM. However, this threshold was predicted to depend on the glycolytic flux (Figures 1C and 3A-B). According to the model, the switch should occur at an acetate concentration of 3.8 mM when the glycolytic activity is 60 % of its maximum value, and at a concentration of just 0.6 mM when the glycolytic activity is reduced to 20 % of the maximum. This suggests that the glycolytic flux determines the role of acetate as a co-substrate or by-product of glycolytic metabolism.

**Figure 3.**
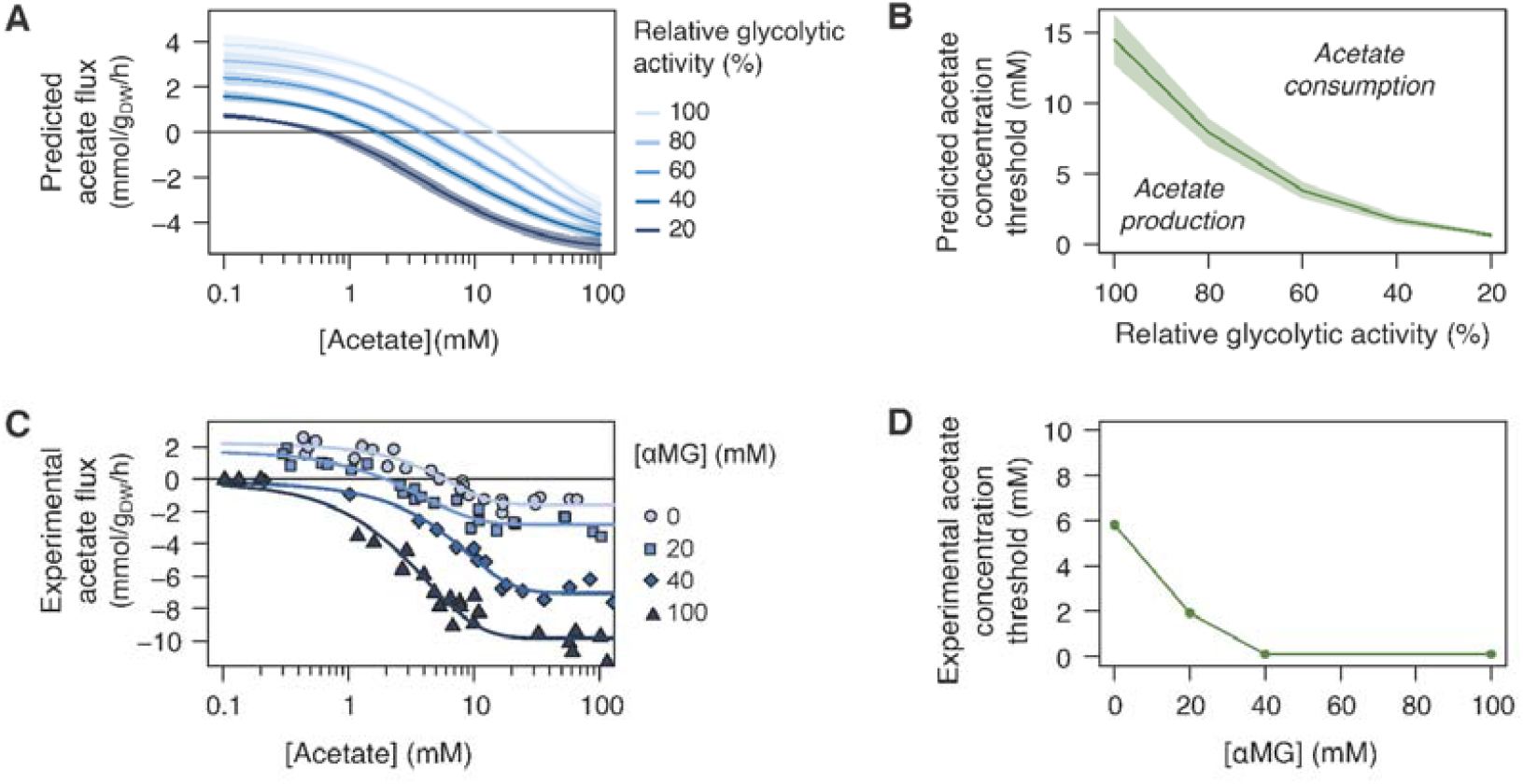
Acetate flux reversal is controlled by the glycolytic flux. A Predicted acetate flux in *E. coli* on glucose (15 mM) as a function of the acetate concentration at different values of glycolytic activity (from 20 to 100 % of the maximal value). Shaded areas correspond to ± one standard deviation on model predictions. B Predicted acetate concentration threshold at which the acetate flux reverses as a function of the glycolytic activity, determined from the data shown in panel (A). Shaded areas correspond to ± one standard deviation on model predictions. C Experimental acetate flux of *E. coli* K-12 MG1655 grown on glucose (15 mM) at different acetate concentrations in the presence of αMG (0, 20, 40 or 100 mM). The data measured in the absence of αMG are reproduced from (Enjalbert *et al*., 2017). Each data point represents an independent biological replicate, and the lines represent best fits using a logistic function. D Experimental acetate concentration threshold at which the acetate flux reverses as a function of the αMG concentration, determined from the best fits shown in panel (C).

We tested this hypothesis by measuring acetate fluxes in *E. coli* grown on glucose (15 mM) plus acetate (from 0.1 to 100 mM) at different concentrations of αMG (0, 20, 40 or 100 mM, Figure 3C). As previously observed at full glycolytic activity (i.e., 0 mM αMG) (Beck *et al*., 2022; Enjalbert *et al*., 2017; Millard *et al*., 2021; Pinhal *et al*., 2019), acetate production decreased when the acetate concentration was increased (Figure 3C), and the acetate flux reversed at a concentration of ∼6 mM (Figure 3C-D), resulting in co-consumption of glucose and acetate. This threshold concentration gradually decreased when the glycolytic flux was reduced (from 2 mM acetate at 20 mM αMG to below 0.1 mM acetate at 40 or 100 mM αMG). Here again, the experimental profiles are qualitatively similar to the simulated profiles, albeit with quantitative differences. These results demonstrate that the acetate concentration threshold at which the acetate flux reverses depends on the glycolytic flux. It therefore follows that when the glycolytic flux is limited, acetate is utilized as a co-substrate at lower acetate concentrations. These results also confirm that inhibiting glycolysis enhances acetate utilization over the full range of acetate concentrations tested here, further validating the unusual control patterns identified above, and generalizing their functional relevance for acetate concentrations spanning three orders of magnitude.

### Acetate enhances *E. coli* growth at low glycolytic flux

Acetate is considered toxic for microbial cells, but the model predicts this might not always be the case. At full glycolytic activity, the model predicts that maximal growth should occur at low acetate concentrations, with the growth rate remaining above 95 % of the maximum only at acetate concentrations below 12 mM (Figures 1D and 4A). In agreement with the reported toxicity of acetate, the growth rate then monotonously decreases for acetate concentrations above 12 mM. However, when the glycolytic flux is reduced, maximal growth is predicted to occur at high acetate concentrations (Figures 1D and 4A). For instance, when the glycolytic activity is reduced to 20 %, the growth rate is expected to be maximal at an acetate concentration of 28 mM, and should remain above 95 % of the maximum over a broad range of acetate concentrations (between 12 and 86 mM). According to the model therefore, acetate should enhance *E. coli* growth at low glycolytic flux.

**Figure 4.**
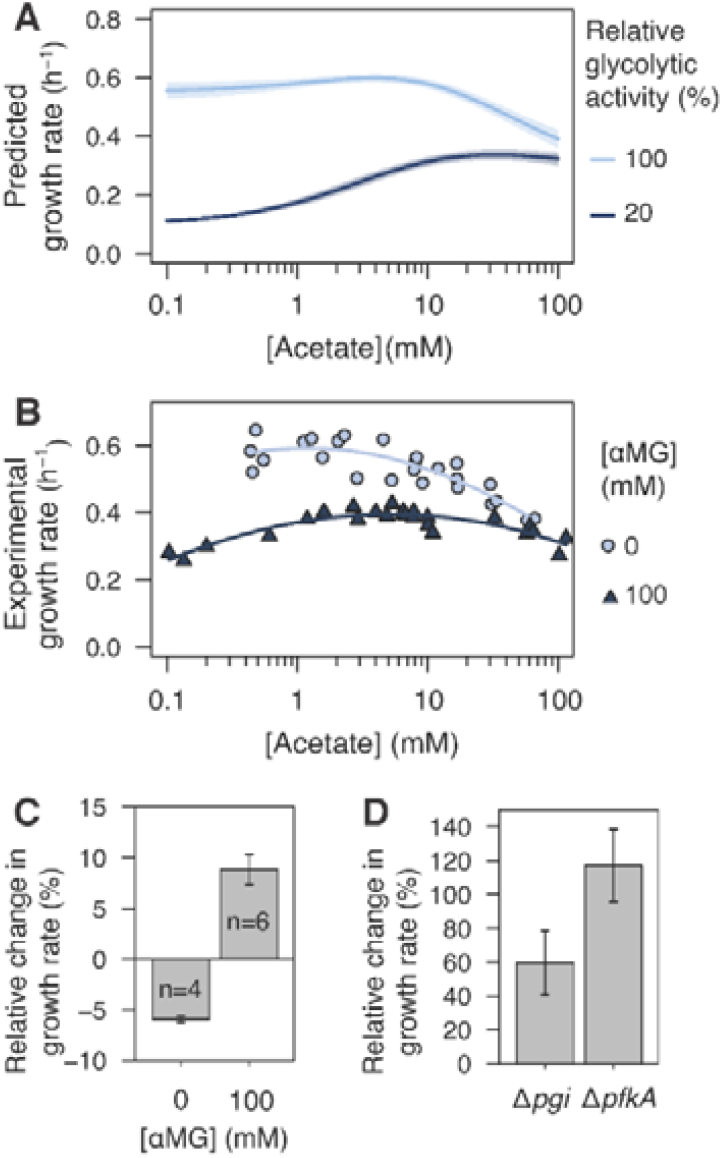
Effect of acetate on *E. coli* growth on glucose. A Predicted growth rate of *E. coli* on glucose (15 mM) as a function of the acetate concentration at 100 and 20 % of the maximal glycolytic activity. Shaded areas correspond to ± one standard deviation. B Experimental growth rate of *E. coli* K-12 MG1655 grown on glucose (15 mM) as a function of the acetate concentration at high (0 mM αMG, light blue, reproduced from (Enjalbert *et al*., 2017)) or low (100 mM αMG, dark blue) glycolytic flux. Each data point represents an independent biological replicate and the lines represent best polynomial fits. C Relative change in the growth rate of *E. coli* K-12 BW25113 wild-type in the presence of 10 mM acetate at high (0 mM αMG) and low (100 mM αMG) glycolytic flux. Mean values ± standard deviations (error bars) were estimated from *n* independent biological replicates, as indicated on the figure. D Relative change in the growth rates of *E. coli* K-12 BW25113 Δ*pgi* and Δ*pfkA* strains in the presence of 10 mM acetate. Mean values ± standard deviations (error bars) were estimated from three independent biological replicates.

To test this prediction, we measured the growth rate of *E. coli* on glucose (15 mM) with acetate (0.1 to 100 mM) in the presence of 100 mM αMG to inhibit glycolysis (Figure 4B). The data measured in the absence of αMG are shown for comparison. At maximal glycolytic flux, the growth rate decreased monotonically with the acetate concentration, but this relationship became non-monotonic when glycolysis was inhibited, which is qualitatively consistent with model predictions. In the presence of αMG, the growth rate was lower in the absence of acetate than at high acetate concentrations (0.28 ± 0.02 h^−1^ with acetate < 0.5 mM versus 0.34 ± 0.01 h^−1^ with 60 mM acetate, p-value = 8.10^−3^). As predicted by the model therefore, acetate appears to be beneficial rather than toxic to *E. coli* when the glycolytic flux is reduced. The growth rate was slightly lower at 100 mM acetate (0.30 ± 0.03 h^−1^) than at 60 mM acetate, but the similar growth rates observed across the concentration range demonstrate that acetate is not toxic when the glycolytic flux is low, even at high concentrations.

As an orthogonal, alternative strategy to the use of a glycolytic inhibitor, we tested the impact of acetate on the growth of mutant strains with low glycolytic flux, namely the *E. coli* K-12 BW25113 Δ*pgi* and Δ*pfkA* strains, where the glycolytic flux is 24 and 29 % that of the wild-type strain, respectively (Long & Antoniewicz, 2019). Their growth rates were measured on 15 mM glucose, with or without 10 mM acetate and the effect of acetate was quantified as the relative change in growth rate. The wild-type isogenic strain was grown in the absence and presence of αMG (100 mM) as a control. As expected, acetate reduced the growth rate of the wild-type strain in the absence of αMG (−6 ± 1 %, p-value = 4.10^−5^) but increased its growth in the presence of 100 mM αMG (+9 ± 2 %, p-value = 3.10^−5^) (Figure 4C). The beneficial effect of acetate on growth was also observed in the Δ*pgi* and Δ*pfkA* strains (Figure 4D), whose growth rate strongly increased despite the absence of αMG (Δ*pgi* : +60 ± 19 %, p-value = 3.10^−2^; Δ*pfkA* : +117 ± 21 %, p-value = 1.10^−2^). These results confirm that the beneficial effect of acetate is not due to the presence of αMG but is a consequence of the decrease in glycolytic flux.

Overall, all results are in line with the predictions of the model and demonstrate that acetate enhances *E. coli* growth when the glycolytic flux is reduced by either chemical or genetic perturbations.

### At low glycolytic flux, acetate activates *E. coli* metabolism and growth at the transcriptional level

At high glycolytic flux, it has been proposed (Millard *et al*., 2021) that the apparent “toxicity” of acetate is due to its inhibition of most metabolic genes – including those for the glucose phosphotransferase system (PTS), glycolysis and TCA cycle. Acetate also increases the expression of genes related to stress responses and downregulates genes related to transport, energy biosynthesis and mobility for instance. To determine if acetate has a similar regulatory effect at low glycolytic flux, we compared the transcriptomes of *E. coli* grown on glucose (15 mM) plus acetate (0, 10, 50 or 100 mM) in the presence of αMG (100 mM), with those measured in the absence of αMG at the same acetate concentrations (Millard *et al*., 2021).

We observed that contrary to what occurs at maximal glycolytic flux (0 mM αMG) acetate did not trigger a stress response at low glycolytic flux (100 mM αMG) (Figure 5A). Conversely, acetate increased gene expression for the respiratory chain and some biosynthetic processes (ribonucleotide monophosphate biosynthesis, amino acid production). Other hallmarks of the effect of acetate at maximal glycolytic flux such as the inhibition of mobility-related gene expression were also not observed at low glycolytic flux. The activation of glutamate catabolism, the repression of transport related genes, and the repression of carbohydrate transport and catabolism were among the few similarities between the responses to acetate at high and low glycolytic flux.

**Figure 5.**
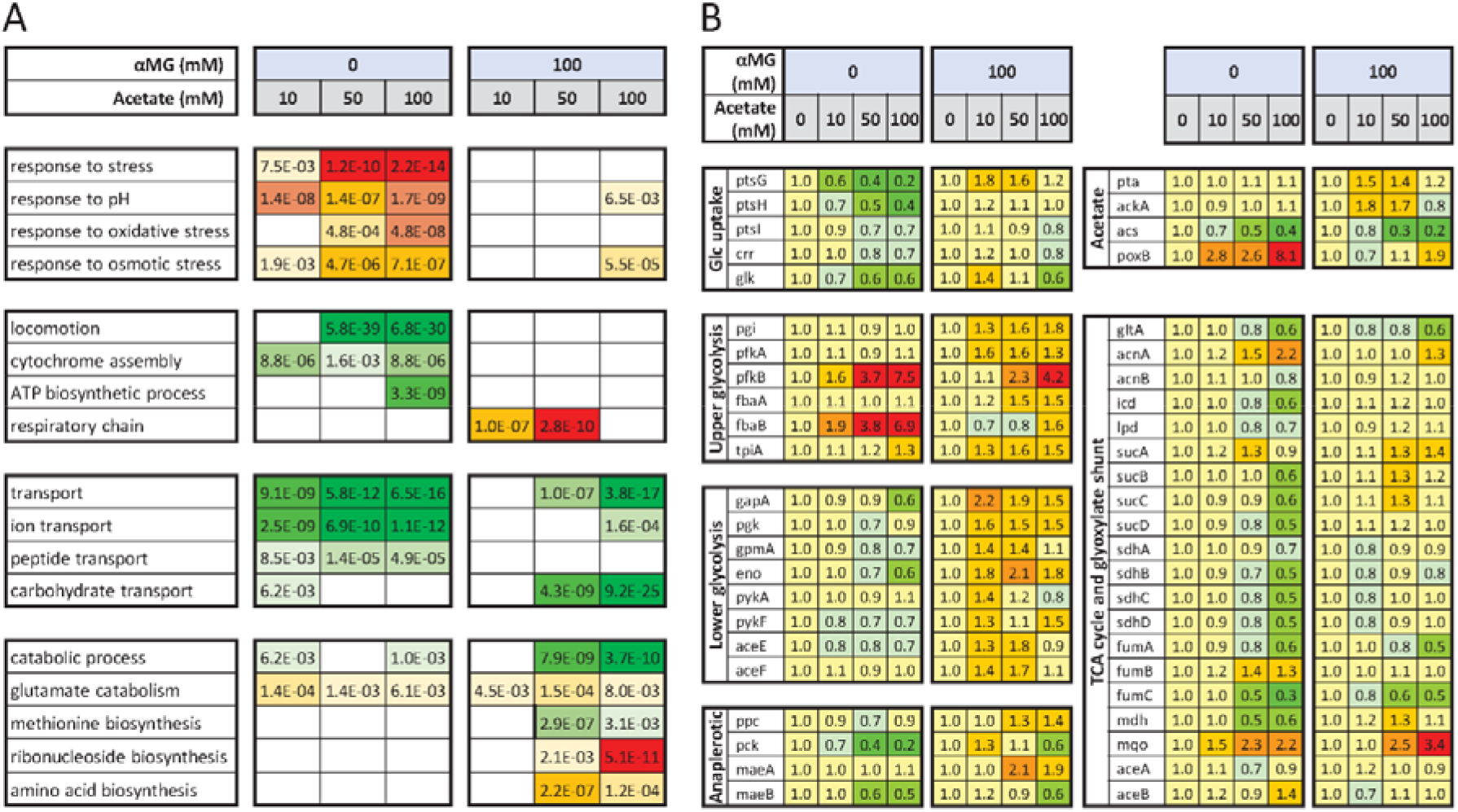
Regulation of *E. coli* transcriptome by acetate at high and low glycolytic flux. A, B Transcriptomic response of *E. coli* K-12 MG1655 to acetate (10, 50 or 100 mM versus 0 mM) at high (0 mM αMG) and low (100 mM αMG) glycolytic flux. The data obtained in the absence of αMG are reproduced from (Millard *et al*., 2021). The biological functions modulated by the presence of acetate (based on Gene Ontology analysis) are shown in part (A), with the corresponding p-values. The expression levels of central metabolic genes are shown in part (B). Gene expression is normalized to the value measured at the same αMG concentration but without acetate. Green and red indicate decreased and increased expression, respectively.

**Figure 6.**
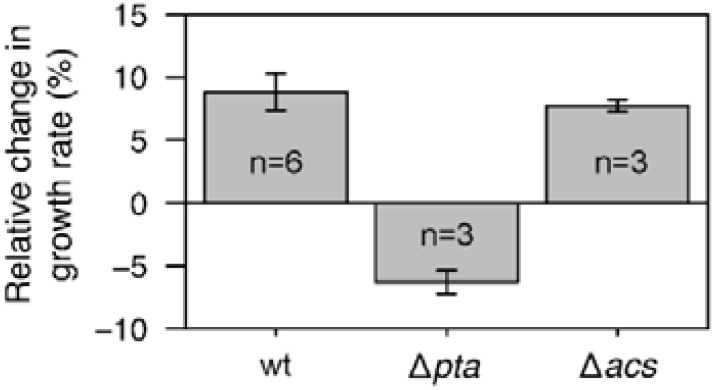
Response of wild-type, Δ*pta*, and Δ*acs* strains of *E. coli* to acetate during growth on glucose. Mean relative change in the growth rates of wild-type, Δ*pta*, and Δ*acs E. coli* K-12 BW25113 strains in the presence of 10 mM acetate at low glycolytic flux (100 mM αMG). Mean values and standard deviations (error bars) were estimated from *n* independent biological replicates, as indicated on the figure.

At the metabolic level, acetate-driven inhibition of the expression of central metabolic genes (PTS, TCA and glycolytic enzymes) and growth machinery (ribosomal proteins) was not observed at low glycolytic flux (Figure 5B). In fact, expression of most of these genes (47/65) was slightly higher in the presence of 50 mM acetate and 100 mM αMG, whereas at maximal glycolytic flux it was lower at these acetate concentrations for a similar proportion (46/65). When the glycolytic flux was inhibited, *ptsG* expression, which controls glucose uptake (Rohwer *et al*, 2000), was 70 % higher at 10 and 50 mM acetate. The expression of most glycolytic genes was 15 to 110 % higher at 50 mM acetate, while the expression of TCA cycle genes remained stable. Regarding acetate metabolism, the expression of *pta* and *ackA* was slightly higher at 10 and 50 mM acetate, while alternative acetate utilization via Acs was inhibited by acetate (5-fold reduction in expression at 100 mM). Finally, expression of genes encoding ribosomal proteins (*rpl, rpm* and *rps* operons) was increased by 57 % on average at 50 mM acetate and 48 % on average at 100 mM acetate (Dataset S1).

These results indicate that the stress response and lower metabolic activity triggered by acetate at high glycolytic flux do not occur at low glycolytic flux. In this situation, acetate enhances the expression of ribosomal and central metabolic genes, in line with its positive effect on growth. This is the opposite of what is observed at high glycolytic flux, demonstrating that the regulatory role of acetate is largely determined by the glycolytic flux.

### Acetate-enhanced growth is supported by the Pta-AckA pathway

According to the model (Figure 1A), the reversible Pta-AckA pathway (acetate consumption or production) explains by itself how acetate promotes *E. coli* growth, without the need for Acs (acetate consumption only). To confirm this hypothesis, we measured the growth rates of *E. coli* K-12 BW25113 wild-type and of Δ*pta* and Δ*acs* isogenic mutant strains cultivated on 15 mM glucose plus 100 mM αMG, with or without 10 mM acetate. The effect of acetate was quantified as the relative change in growth rate induced. Acetate had a similar, beneficial effect on the growth of the wild-type strain (+9 ± 2 %, p-value = 3.10^−5^) and of the Δ*acs* strain (+8 ± 1 %, p-value = 1.10^−3^), but not for the Δ*pta* strain, whose growth rate decreased slightly but significantly (−6 ± 1 %, p-value = 7.10^−3^). These results confirm that the beneficial effect of acetate on *E. coli* growth at low glycolytic flux is mediated by the Pta-AckA pathway, with no involvement of Acs in this phenomenon.

### Acetate boosts growth of *E. coli* on glycerol and galactose

Assuming that the beneficial impact of acetate on *E. coli* growth stems from the tight interplay of glycolytic and acetate fluxes, this should be independent of the nature of the carbon source, implying that acetate should boost *E. coli* growth on glycolytic substrates with low glycolytic flux, such as glycerol and galactose (Gerosa *et al*, 2015). And indeed, wild-type *E. coli* growth was enhanced on both glycerol (+7 ± 1 %, p-value = 1.10^−2^) and galactose (+52 ± 1 %, p-value = 2.10^−4^) in the presence of 10 mM acetate (Figure 7). Similar levels of enhancement were observed for the Δ*acs* strain (+8 ± 1 % on glycerol, p-value = 3.10^−4^; +53 ± 5 % on galactose, p-value = 3.10^−3^), but not for the Δ*pta* strain (−8 ± 2 % on glycerol, p-value = 1.10^−2^; +2 ± 12 % on galactose, p-value = 0.70). These results demonstrate that acetate promotes *E. coli* growth on galactose and glycerol. As observed for glucose, the growth enhancement is mediated exclusively by the Pta-AckA pathway, with no involvement of Acs. These results extend and generalize our findings to glycolytic carbon sources other than glucose and support our findings in more physiological conditions, i.e. without an inhibitor or deletion of metabolic genes.

**Figure 7.**
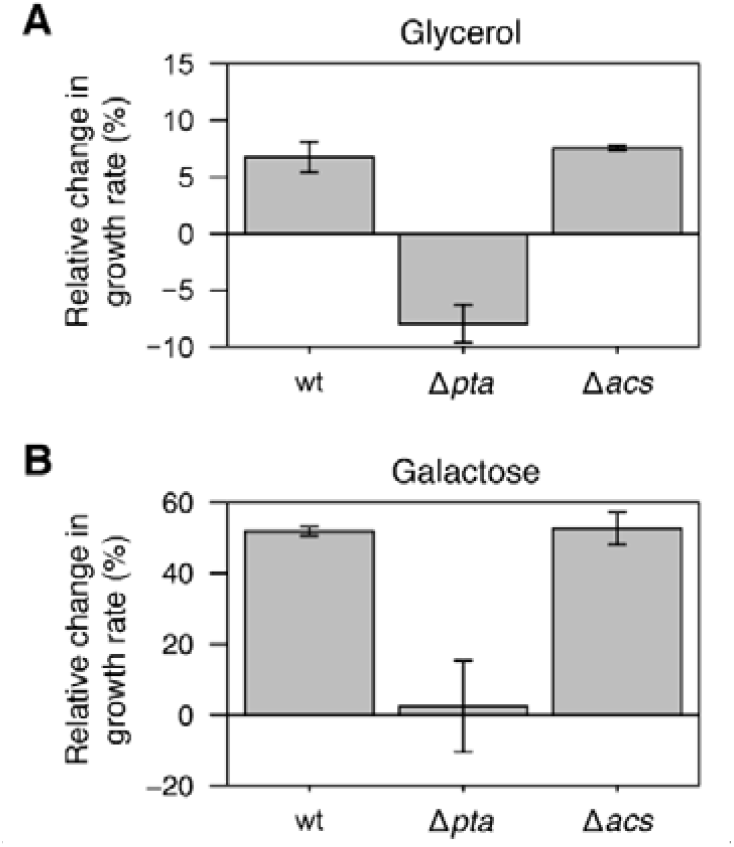
Response of *E. coli* to acetate during growth on glycerol or galactose. A,B Impact of the presence of 10 mM acetate on the growth rate of wild-type, Δ*pta*, and Δ*acs E. coli* K-12 BW25113 strains grown on (A) 30 mM glycerol and (B) 15 mM galactose. Mean values and standard deviations (error bars) were estimated from three independent biological replicates.

## Discussion

Following a systems biology approach, we used a kinetic model of *E. coli* metabolism to generate a comprehensive map of the functional relationship between glycolytic and acetate metabolisms and predict their combined impact on *E. coli* growth. In agreement with the predictions of the model, experimental results revealed a strong interplay between acetate and glycolytic fluxes in *E. coli*, with wide-ranging physiological consequences for the cells. Despite its established toxicity for microbial growth, we found that acetate improves the robustness of *E. coli* to glycolytic perturbations, and even boosts growth when the glycolytic flux is low.

Previous studies of the glycolytic control of acetate metabolism were carried out in the absence of acetate. In this situation, as observed here, glycolysis as a positive effect on acetate production. However, our results also reveal that in the presence of acetate, glycolysis negatively affects acetate utilization. Glycolytic control of the acetate flux is thus determined by the concentration of acetate itself. Because of this non-monotonous control pattern, reduced glycolytic flux increases acetate uptake and lowers the acetate concentration threshold at which acetate is co-consumed with glucose, with the overall effect of increasing acetate utilization by *E. coli*. This demonstrates that the metabolic status of acetate as a by-product or co-substrate of glycolytic carbon sources is determined both by the concentration of acetate (Enjalbert *et al*., 2017; Millard *et al*., 2021) and the glycolytic flux, and thus by the availability of glycolytic nutrients. This mechanism is exclusively mediated by the Pta-AckA pathway and its role and activity are determined by the nutritional conditions experienced by *E. coli*. The Pta-AckA pathway can therefore be seen as a metabolic valve that integrates information on the nutritional environment of the cells to rapidly adjust the acetate flux depending on the availability of glycolytic substrates and acetate. Our results indicate that these properties, which emerge simply from the distribution of control throughout the network, buffers the total carbon uptake flux and makes *E. coli* more robust to changes in the nutritional environment.

When the glycolytic flux is low, enhanced acetate utilization promotes *E. coli* growth, demonstrating that acetate can be beneficial to *E. coli*, even at high concentrations. Whereas 60 mM acetate has been shown to inhibit growth by ∼35 % at maximal glucose uptake rates (Enjalbert *et al*., 2017; Luli & Strohl, 1990; Pinhal *et al*., 2019), the same concentration of acetate had a positive effect on growth at low glucose uptake rates (+21 % in presence of 100 mM αMG). We confirmed the beneficial effect of acetate on *E. coli* growth using three orthogonal strategies to reduce the glycolytic flux: i) chemical inhibition of glucose uptake with αMG, ii) mutant strains deleted for glycolytic genes, and ii) glycolytic substrates with a naturally low glycolytic flux (glycerol and galactose). Since the impact of acetate on growth is partly determined by the glycolytic flux, this metabolic by-product is not toxic *per se*. Importantly, these results also weaken several hypotheses that have been proposed to explain the apparent “toxicity” of acetate for microbial growth. Indeed, perturbations of the proton gradient, of the anion composition of the cell, or of methionine biosynthesis, would have a monotonic impact on growth. The slight but global activating effect of acetate on the transcriptome at low glycolytic flux, which correlates with the growth response, also contrasts with its inhibitory effect at high glycolytic flux. The results of this study indicate that mechanisms that only explain growth inhibition at high glycolytic flux are inadequate. Likewise, the recent suggestion that gluconeogenic substrates such as acetate enhance *E. coli* growth on glycolytic substrates (as observed for mixtures of oxaloacetate with glycerol, xylose or fucose) (Okano *et al*, 2020) also fails to capture the non-monotonic growth response of *E. coli* to acetate. The regulatory effect of acetate on *E. coli* metabolism, which depends both on the acetate concentration and on the glycolytic flux, is therefore more complex than previously suggested (Millard *et al*., 2021) and should be investigated further.

Previous reports have shown that the “toxicity” of acetate for *E. coli* growth depends on the pH (Lawford & Rousseau, 1993; Orr *et al*, 2019; Pinhal *et al*., 2019). In our experiments, the pH was initially set to 7.0 but was not adjusted during growth. Nevertheless, the high buffer capacity of the M9 cultivation medium ensured that the pH remained stable in the exponential growth phase where growth rates and extracellular fluxes were measured, with changes of less than 0.3 pH unit in all experiments (Figure S1). Therefore, the metabolic and physiological responses of *E. coli* to acetate highlighted in this study are not due to changes in pH. Another potential issue is that the effect of acetate might also depend on the concentration of salts (Bae & Lee, 2017; Lee *et al*, 2010). The effect of acetate added as sodium salt may differ from its effect when added as acetic acid. However, repeat experiments performed with acetic acid instead of sodium acetate showed that the beneficial effect of acetate on *E. coli* growth on glucose, galactose and glycerol was maintained (Figure S2).

The measured responses of *E. coli* to a broad range of perturbations of the glycolytic and acetate metabolisms were qualitatively consistent with model predictions, though quantitative differences were observed. These differences may stem from the data used to construct and calibrate the model, which were obtained exclusively at maximal glycolytic flux. Therefore, although it was not initially developed to address the questions of this study, the kinetic model used here successfully predicts the observed interplay between glycolytic and acetate metabolisms, as well as the beneficial effect of acetate on *E. coli* growth at low glycolytic flux. Since the model does not account for the activation by acetate of the expression of glucose uptake, glycolytic, and ribosomal genes, this suggests that transcriptional activation by acetate may not be crucial for the flux response of *E. coli*. The metabolic network may indeed directly sense and integrate the availability of glucose and acetate to coordinate glucose and acetate fluxes, as already reported for the coordination other metabolic processes (Doucette *et al*, 2011; Millard *et al*, 2017; Yuan *et al*, 2009). In keeping with this hypothesis, our results support the concept of acetate flux being thermodynamically controlled by acetate itself, as detailed in (Enjalbert *et al*., 2017), and we show that the thermodynamically-driven flux reversal is also controlled by the glycolytic flux. The reduction in glucose-derived acetyl-CoA is counterbalanced by an increased contribution of acetate, which sustains growth. Metabolic regulation is thus (at least partly) responsible for the interplay between acetate and glycolytic flux. The precise regulatory program driven by acetate remains to be identified.

Acetate concentrations in the gut range from 30 to 100 mM and glycolytic nutrients, such as glycerol and galactose (Fabich *et al*., 2008; Matamouros *et al*, 2018; Snider *et al*, 2009), are generally scarce. Given the established inhibition of microbial growth by acetate, these high acetate levels were considered detrimental to *E. coli* colonization. Our results indicate that acetate, far from being toxic, may in fact be beneficial to *E. coli* in this environment. Acetate enhances *E. coli* growth when glycolytic substrates are limiting and may therefore facilitate its colonization and survival in the gut. Moreover, acetate stabilizes growth when the glycolytic flux is perturbed by temporary nutrient shortages, which are frequent in the gut. We suggest that the underlying regulatory mechanism may therefore be optimal for *E. coli* in its natural environment. Our results also call for the role of acetate in the gut to be reconsidered, since it represents a valuable and abundant resource that may benefit *E. coli* and other microorganisms.

In biotechnology, developing efficient strategies to overcome the negative effects of acetate requires understanding what controls the acetate flux and how in turn acetate controls microbial metabolism. By addressing these two needs, our findings should immediately help design bioprocesses with reduced acetate overflow and accelerate the development of mixed feed processes in which acetate is utilized alongside glucose or other glycolytic substrates. Current strategies are mainly based on increasing Acs activity to (re)use acetate present in the feedstock, provided as a co-substrate or produced by the Pta-AckA pathway. Given the Pta-AckA pathway produces one ATP and Acs consumes two ATPs, strategies based on activating the Pta-AckA-Acs cycle are highly energy-consuming (Valgepea *et al*., 2010). Strategies involving the Pta-AckA pathway alone are inherently less costly for the cell. Our results show that the Pta-AckA pathway enables efficient co-utilization of acetate and glycolytic substrates (glucose, glycerol, galactose) in *E. coli* by itself, with a positive effect on growth. The comprehensive understanding of the relationship between glycolytic and acetate metabolisms provided by this study may also represents a valuable guide to increase the productivity of valuable products synthesized from acetate-derived AcCoA (Becker & Wittmann, 2015; Zhang *et al*, 2015).

*E. coli* is not the only microorganism that co-consumes glucose with fermentation by-products. *Saccharomyces cerevisiae* can co-consume ethanol and glucose (Raamsdonk *et al*, 2001; Xiao *et al*, 2022), and mammalian cells co-consume lactate with glucose (Faubert *et al*, 2017; Hui *et al*, 2020; Hui *et al*, 2017; Rabinowitz & Enerbäck, 2020). Our findings may thus be generalizable to other microorganisms. Glycolytic control of fermentative pathways has indeed been observed, albeit poorly understood, in virtually all microorganisms. Our results explain why co-consumption of ethanol with glucose in yeast is observed at low glucose uptake (Raamsdonk *et al*., 2001). A similar relationship between glycolytic and lactate fluxes has also been reported *in vitro* in mammalian cells (Mulukutla *et al*, 2012), with the switch between lactate production and utilization being determined by the glycolytic flux. It has been suggested that lactate rather than glucose is the primary circulating energy source in the human body (Faubert *et al*., 2017; Rabinowitz & Enerbäck, 2020). Our study demonstrates that similarly to lactate, acetate is not a deleterious waste product but a valuable nutrient that can be shuttled between cells. The reversible Pta-AckA pathway enables uncoupling of glycolysis from the TCA cycle, indicating that this evolutionarily conserved design principle of eukaryotic metabolism (Xiao *et al*., 2022) is also present in bacteria. As a nutrient and global regulator of microbial metabolism, acetate plays a major role in cellular, microbial community and host-level carbon and energy homeostasis.

## Materials and Methods

### Strains

*Escherichia coli* K-12 MG1655 was chosen as the model wild-type strain. For experiments involving mutant strains, *E. coli* K-12 BW25113 was chosen as the reference wild-type strain, and its isogenic Δ*pta*, Δ*pgi* and Δ*pfkA* mutants were constructed from the KEIO collection (Baba *et al*, 2006) by removing the kanamycin cassette (Datsenko & Wanner, 2000). The Δ*acs* strain was constructed by CRISPR-Cas9 genome editing using the method and plasmids described in Wei et al. (Jiang *et al*, 2015). *pta* deletion was confirmed by genome sequencing and *acs, pgi* and *pfkA* deletions by PCR (with primers listed in Table S1).

### Growth conditions

*E. coli* was grown in M9 minimal media (Millard *et al*, 2014) supplemented with 15 mM glucose, 15 mM galactose, or 30 mM glycerol. Sodium acetate (prepared as a 1.2 M solution at pH 7.0 by addition of potassium hydroxide), acetic acid (prepared as a 1.2 M solution at pH 7.0 by addition of ammonium hydroxide) and methyl α-D-glucopyranoside (αMG, prepared as a 1.2 M solution) were added up to the required concentrations. The cells were grown in shake flasks at 37 °C and 200 rpm, in 50 mL of medium. Growth was monitored by measuring the optical density (OD) at 600 nm using a Genesys 6 spectrophotometer (Thermo, USA). Biomass concentrations were determined using a conversion factor of 0.37 g_DW_/L/OD unit (Revelles *et al*, 2013).

### Transcriptomics experiments

Cells were grown in M9 medium with 15 mM glucose, 100 mM αMG, and 0, 10, 50, or 100 mM sodium acetate. Sample preparations and transcriptomics analyses were carried out as described previously (Millard *et al*., 2021). Three independent replicates were analyzed for each condition. The transcriptomics data can be downloaded from the ArrayExpress database (www.ebi.ac.uk/arrayexpress) under accession number E-MTAB-11717. Transcriptomics data in the absence of glycolytic inhibition (ArrayExpress database accession number E-MTAB-9086 (Millard *et al*., 2021)) were measured under the exact same conditions but without αMG.

### Metabolomics experiments

Extracellular concentrations of glucose and acetate were quantified in 180 µL of filtered broth (0.2 μm syringe filter, Sartorius, Germany) by 1D ^1^H-NMR on a Bruker Avance 500 MHz spectrometer equipped with a 5-mm z-gradient BBI probe (Bruker, Germany), as described previously (Millard *et al*., 2021).

### Flux calculation

Glucose uptake, acetate and growth rates were calculated from glucose, acetate and biomass concentration–time profiles using PhysioFit (v1.0.2, https://github.com/MetaSys-LISBP/PhysioFit) (Peiro *et al*, 2019).

### Kinetic modeling

We used a kinetic model of *E. coli* (Millard *et al*., 2021) available from the Biomodels database (https://www.ebi.ac.uk/biomodels) (Chelliah *et al*, 2015) under identifier MODEL2005050001. Simulations were carried out as described in the main text, using COPASI (Hoops *et al*, 2006) (v4.34) with the CoRC package (Forster *et al*, 2020) (COPASI R Connector v0.11, https://github.com/jpahle/CoRC) in R (v4.1.3, https://www.r-project.org). The R notebook used to perform the simulations and to generate the figures is provided at https://github.com/MetaSys-LISBP/glucose_acetate_interplay.

### Sensitivity analyses

We used a Monte-Carlo approach (Saa & Nielsen, 2017) to estimate the robustness of model predictions. Following the procedure detailed in (Millard *et al*., 2021), 500 simulated sets of calibration data were generated with noise added according to experimental standard deviations. For each of these artificially noisy data sets, we estimated model parameters and ran all simulations from the optimal parameters values. We calculated the mean value and standard deviation of each flux from the distribution of values obtained for the 500 sets of parameters.

### Statistical analyses

Comparisons between strains or conditions were performed using Student’s t-tests with two-tailed distributions. The R notebook used to perform the calculations is provided at https://github.com/MetaSys-LISBP/glucose_acetate_interplay.

## Data Availability

The data used to make the figures can be found in the Source Data files. The data, model and computer code produced in this study are available in the following databases:

- Transcriptomics data : ArrayExpress E-MTAB-11717 (https://www.ebi.ac.uk/biostudies/arrayexpress/studies/E-MTAB-11717)
- Model: BioModels MODEL2005050001 (https://www.ebi.ac.uk/biomodels/MODEL2005050001)
- Code: GitHub (https://github.com/MetaSys-LISBP/glucose_acetate_interplay)

## Acknowledgments

This work was supported by a grant to Pierre Millard and Thomas Gosselin-Monplaisir from the Département MICA of INRAE and the Région Occitanie (grant COCA-COLI) and from the Toulouse Biotechnology Institute (grant Master-2022). The authors thank MetaboHub-MetaToul (Metabolomics and Fluxomics facilities, Toulouse, France, http://www.metatoul.fr), part of the French National Infrastructure for Metabolomics and Fluxomics (http://www.metabohub.fr) funded by the ANR (MetaboHUB-ANR-11-INBS-0010), for access to NMR facilities. The authors are grateful to Edith Vidal and Abdulrahman Khabbaz for help with the growth experiments, and the following INSA Toulouse students for help with the growth and transcriptomics experiments: Lola Blayac, Arno Bruel, Alexis Charbinat, Romane Ducloux, Justine Dumas-Perdriau, Solène Frapard, Sophie Germain, Benjamin Jung, Oliver Larrousse, Nicolas Papadopoulos, Marine Rodeghiero, Leyre Sarrias and Auriane Thomas. The authors also thank the Département Génie Biologique of INSA Toulouse for ongoing support, and Muriel Cocaign-Bousquet and Guy Lippens for insightful discussions on the manuscript.

## Author contributions

**Pierre Millard:** Conceptualization; data curation; formal analysis; funding acquisition; investigation; methodology; project administration; resources; software; supervision; validation; visualization; writing – original draft; writing – review and editing. **Thomas Gosselin-Monplaisir:** Data curation; formal analysis; investigation; methodology; resources; validation; writing – review and editing. **Sandrine Uttenweiler-Joseph:** Methodology; resources; writing – review and editing. **Brice Enjalbert:** Data curation; formal analysis; investigation; methodology; resources; supervision; validation; visualization; writing – review and editing.

## Disclosure and competing interests statement

The authors declare that they have no conflict of interest.

## Notes

https://github.com/MetaSys-LISBP/glucose_acetate_interplay

### Competing Interest Statement

The authors have declared no competing interest.

### Summary of Updates

On the modeling side, we have tested the robustness of the predictions by performing a global sensitivity analysis and a detailed comparison of the numerical predictions with experimental data. On the experimental side, we have carried out new experiments that demonstrate that the beneficial effect of acetate is not due to changes in pH and is conserved when using acetic acid instead of sodium acetate. We have also constructed mutant strains with reduced glycolytic flux and analyzed their response to acetate. These new results i) confirm that the model and its predictions are robust to parameter uncertainty, ii) highlight the consistency of the model predictions with experimental data, iii) rule out a potential role of pH and salts in the observed phenotypes and iv) strengthen our findings in terms of the fundamental principles governing the interplay between glycolytic and acetate metabolisms. These results fully support the conceptual advance provided in the first version.

## References

Baba T, Ara T, Hasegawa M, Takai Y, Okumura Y, Baba M, Datsenko KA, Tomita M, Wanner BL, Mori H (2006) Construction of Escherichia coli K-12 in-frame, single-gene knockout mutants: the Keio collection. Mol Syst Biol 2: 2006

Bae YM, Lee SY (2017) Effect of salt addition on acid resistance response of Escherichia coli O157:H7 against acetic acid. Food Microbiol 65: 74–82

Beck AE, Pintar K, Schepens D, Schrammeck A, Johnson T, Bleem A, Du M, Harcombe WR, Bernstein HC, Heys JJ et al (2022) Environment constrains fitness advantages of division of labor in microbial consortia engineered for metabolite push or pull interactions. mSystems: e0005122

Becker J, Wittmann C (2015) Advanced biotechnology: metabolically engineered cells for the bio-based production of chemicals and fuels, materials, and health-care products. Angew Chem Int Ed Engl 54: 3328–3350

Bernal V, Castano-Cerezo S, Canovas M (2016) Acetate metabolism regulation in Escherichia coli: carbon overflow, pathogenicity, and beyond. Appl Microbiol Biotechnol 100: 8985–9001

Bren A, Park JO, Towbin BD, Dekel E, Rabinowitz JD, Alon U (2016) Glucose becomes one of the worst carbon sources for E. coli on poor nitrogen sources due to suboptimal levels of cAMP. Scientific reports 6: 24834

Chelliah V, Juty N, Ajmera I, Ali R, Dumousseau M, Glont M, Hucka M, Jalowicki G, Keating S, Knight-Schrijver V et al (2015) BioModels: ten-year anniversary. Nucleic Acids Res 43: D542–548

Cummings JH, Englyst HN (1987) Fermentation in the human large intestine and the available substrates. Am J Clin Nutr 45: 1243–1255

Cummings JH, Macfarlane GT, Englyst HN (2001) Prebiotic digestion and fermentation. Am J Clin Nutr 73: 415S–420S

Datsenko KA, Wanner BL (2000) One-step inactivation of chromosomal genes in Escherichia coli K-12 using PCR products. Proc Natl Acad Sci U S A 97: 6640–6645

de Graaf AA, Maathuis A, de Waard P, Deutz NE, Dijkema C, de Vos WM, Venema K (2010) Profiling human gut bacterial metabolism and its kinetics using [U-13C]glucose and NMR. NMR Biomed 23: 2–12

De Mey M, De Maeseneire S, Soetaert W, Vandamme E (2007) Minimizing acetate formation in E. coli fermentations. J Ind Microbiol Biotechnol 34: 689–700

Doucette CD, Schwab DJ, Wingreen NS, Rabinowitz JD (2011) alpha-Ketoglutarate coordinates carbon and nitrogen utilization via enzyme I inhibition. Nat Chem Biol 7: 894–901

Eiteman MA, Altman E (2006) Overcoming acetate in Escherichia coli recombinant protein fermentations. Trends Biotechnol 24: 530–536

Enjalbert B, Millard P, Dinclaux M, Portais JC, Letisse F (2017) Acetate fluxes in Escherichia coli are determined by the thermodynamic control of the Pta-AckA pathway. Scientific reports 7: 42135

Fabich AJ, Jones SA, Chowdhury FZ, Cernosek A, Anderson A, Smalley D, McHargue JW, Hightower GA, Smith JT, Autieri SM et al (2008) Comparison of carbon nutrition for pathogenic and commensal Escherichia coli strains in the mouse intestine. Infect Immun 76: 1143–1152

Faubert B, Li KY, Cai L, Hensley CT, Kim J, Zacharias LG, Yang C, Do QN, Doucette S, Burguete D et al (2017) Lactate metabolism in human lung tumors. Cell 171: 358–371 e359

Forster J, Bergmann FT, Pahle J (2020) CoRC - the COPASI R Connector. Bioinformatics

Fuentes LG, Lara AR, Martinez LM, Ramirez OT, Martinez A, Bolivar F, Gosset G (2013) Modification of glucose import capacity in Escherichia coli: physiologic consequences and utility for improving DNA vaccine production. Microb Cell Fact 12: 42

Gerosa L, Haverkorn van Rijsewijk BR, Christodoulou D, Kochanowski K, Schmidt TS, Noor E, Sauer U (2015) Pseudo-transition analysis identifies the key regulators of dynamic metabolic adaptations from steady-state data. Cell systems 1: 270–282

Harden A (1901) The chemical action of Bacillus coli communis and similar organisms on carbohydrates and allied compounds. J Chem Soc Trans 79: 610–628

Hollowell CA, Wolin MJ (1965) Basis for the exclusion of Escherichia coli from the rumen ecosystem. Appl Microbiol 13: 918–924

Hoops S, Sahle S, Gauges R, Lee C, Pahle J, Simus N, Singhal M, Xu L, Mendes P, Kummer U (2006) COPASI--a COmplex PAthway SImulator. Bioinformatics 22: 3067–3074

Hui S, Cowan AJ, Zeng X, Yang L, TeSlaa T, Li X, Bartman C, Zhang Z, Jang C, Wang L et al (2020) Quantitative fluxomics of circulating metabolites. Cell Metab 32: 676–688 e674

Hui S, Ghergurovich JM, Morscher RJ, Jang C, Teng X, Lu W, Esparza LA, Reya T, Le Z, Yanxiang Guo J et al (2017) Glucose feeds the TCA cycle via circulating lactate. Nature 551: 115–118

Jiang Y, Chen B, Duan C, Sun B, Yang J, Yang S (2015) Multigene editing in the Escherichia coli genome via the CRISPR-Cas9 system. Appl Environ Microbiol 81: 2506–2514

Jones SA, Chowdhury FZ, Fabich AJ, Anderson A, Schreiner DM, House AL, Autieri SM, Leatham MP, Lins JJ, Jorgensen M et al (2007) Respiration of Escherichia coli in the mouse intestine. Infect Immun 75: 4891–4899

Kleman GL, Strohl WR (1994) Acetate metabolism by Escherichia coli in high-cell-density fermentation. Appl Environ Microbiol 60: 3952–3958

Klinke HB, Thomsen AB, Ahring BK (2004) Inhibition of ethanol-producing yeast and bacteria by degradation products produced during pre-treatment of biomass. Appl Microbiol Biotechnol 66: 10–26

Krivoruchko A, Zhang Y, Siewers V, Chen Y, Nielsen J (2015) Microbial acetyl-CoA metabolism and metabolic engineering. Metab Eng 28: 28–42

Kutscha R, Pflügl S (2020) Microbial upgrading of acetate into value-added products— Examining microbial diversity, bioenergetic constraints and metabolic engineering approaches. International journal of molecular sciences 21: 8777

Lawford HG, Rousseau JD (1993) Effects of pH and acetic acid on glucose and xylose metabolism by a genetically engineered ethanologenic Escherichia coli. Applied biochemistry and biotechnology 39-40: 301–322

Lee SY, Rhee MS, Dougherty RH, Kang DH (2010) Antagonistic effect of acetic acid and salt for inactivating Escherichia coli O157:H7 in cucumber puree. J Appl Microbiol 108: 1361–1368

Long CP, Antoniewicz MR (2019) Metabolic flux responses to deletion of 20 core enzymes reveal flexibility and limits of E. coli metabolism. Metab Eng 55: 249–257

Luli GW, Strohl WR (1990) Comparison of growth, acetate production, and acetate inhibition of Escherichia coli strains in batch and fed-batch fermentations. Appl Environ Microbiol 56: 1004–1011

Macfarlane GT, Gibson GR, Cummings JH (1992) Comparison of fermentation reactions in different regions of the human colon. J Appl Bacteriol 72: 57–64

Matamouros S, Hayden HS, Hager KR, Brittnacher MJ, Lachance K, Weiss EJ, Pope CE, Imhaus AF, McNally CP, Borenstein E et al (2018) Adaptation of commensal proliferating Escherichia coli to the intestinal tract of young children with cystic fibrosis. Proc Natl Acad Sci U S A 115: 1605–1610

Millard P, Enjalbert B, Uttenweiler-Joseph S, Portais JC, Letisse F (2021) Control and regulation of acetate overflow in Escherichia coli. Elife 10

Millard P, Massou S, Wittmann C, Portais JC, Letisse F (2014) Sampling of intracellular metabolites for stationary and non-stationary 13C metabolic flux analysis in Escherichia coli. Anal Biochem 465: 38–49

Millard P, Smallbone K, Mendes P (2017) Metabolic regulation is sufficient for global and robust coordination of glucose uptake, catabolism, energy production and growth in Escherichia coli. PLoS Comput Biol 13: e1005396

Mulukutla BC, Gramer M, Hu WS (2012) On metabolic shift to lactate consumption in fedbatch culture of mammalian cells. Metab Eng 14: 138–149

Nanchen A, Schicker A, Sauer U (2006) Nonlinear dependency of intracellular fluxes on growth rate in miniaturized continuous cultures of Escherichia coli. Appl Environ Microbiol 72: 1164–1172

Negrete A, Ng WI, Shiloach J (2010) Glucose uptake regulation in E. coli by the small RNA SgrS: comparative analysis of E. coli K-12 (JM109 and MG1655) and E. coli B (BL21). Microb Cell Fact 9: 75

Okano H, Hermsen R, Kochanowski K, Hwa T (2020) Regulation underlying hierarchical and simultaneous utilization of carbon substrates by flux sensors in Escherichia coli. Nature microbiology 5: 206–215

Orr JS, Christensen DG, Wolfe AJ, Rao CV (2019) Extracellular acidic pH inhibits acetate consumption by decreasing gene transcription of the tricarboxylic acid cycle and the glyoxylate shunt. J Bacteriol 201

Palmqvist E, Hahn-Hägerdal B (2000) Fermentation of lignocellulosic hydrolysates. I: inhibition and detoxification. Bioresource Technology 74

Pasteur L (1857) Mémoire sur la fermentation alcoolique. Comptes Rendus Séances de l’Academie des Sciences 45: 913–916

Peebo K, Valgepea K, Nahku R, Riis G, Oun M, Adamberg K, Vilu R (2014) Coordinated activation of PTA-ACS and TCA cycles strongly reduces overflow metabolism of acetate in Escherichia coli. Appl Microbiol Biotechnol 98: 5131–5143

Peiro C, Millard P, de Simone A, Cahoreau E, Peyriga L, Enjalbert B, Heux S (2019) Chemical and metabolic controls on dihydroxyacetone metabolism lead to suboptimal growth of Escherichia coli. Appl Environ Microbiol 85

Pinhal S, Ropers D, Geiselmann J, de Jong H (2019) Acetate metabolism and the inhibition of bacterial growth by acetate. J Bacteriol 201

Raamsdonk LM, Diderich JA, Kuiper A, van Gaalen M, Kruckeberg AL, Berden JA, Van Dam K (2001) Co-consumption of sugars or ethanol and glucose in a Saccharomyces cerevisiae strain deleted in the HXK2 gene. Yeast 18: 1023–1033

Rabinowitz JD, Enerbäck S (2020) Lactate: the ugly duckling of energy metabolism. Nature Metabolism 2: 566–571

Renilla S, Bernal V, Fuhrer T, Castano-Cerezo S, Pastor JM, Iborra JL, Sauer U, Canovas M (2012) Acetate scavenging activity in Escherichia coli: interplay of acetyl-CoA synthetase and the PEP-glyoxylate cycle in chemostat cultures. Appl Microbiol Biotechnol 93: 2109–2124

Revelles O, Millard P, Nougayrede JP, Dobrindt U, Oswald E, Letisse F, Portais JC (2013) The carbon storage regulator (Csr) system exerts a nutrient-specific control over central metabolism in Escherichia coli strain Nissle 1917. PLoS One 8: e66386

Roe AJ, McLaggan D, Davidson I, O’Byrne C, Booth IR (1998) Perturbation of anion balance during inhibition of growth of Escherichia coli by weak acids. J Bacteriol 180: 767–772

Roe AJ, O’Byrne C, McLaggan D, Booth IR (2002) Inhibition of Escherichia coli growth by acetic acid: a problem with methionine biosynthesis and homocysteine toxicity. Microbiology (Reading) 148: 2215–2222

Rohwer JM, Meadow ND, Roseman S, Westerhoff HV, Postma PW (2000) Understanding glucose transport by the bacterial phosphoenolpyruvate:glycose phosphotransferase system on the basis of kinetic measurements in vitro. J Biol Chem 275: 34909–34921

Russel JB (1992) Another explanation for the toxicity of fermentation acids at low pH: anion accumulation versus uncoupling. J Appl Bacteriol 73: 363–370

Saa PA, Nielsen LK (2017) Formulation, construction and analysis of kinetic models of metabolism: A review of modelling frameworks. Biotechnol Adv 35: 981–1003

Shen Q, Zhao L, Tuohy KM (2011) High-level dietary fibre up-regulates colonic fermentation and relative abundance of saccharolytic bacteria within the human faecal microbiota in vitro. European journal of nutrition

Snider TA, Fabich AJ, Conway T, Clinkenbeard KD (2009) E. coli O157:H7 catabolism of intestinal mucin-derived carbohydrates and colonization. Vet Microbiol 136: 150–154

Sun L, Lee JW, Yook S, Lane S, Sun Z, Kim SR, Jin YS (2021) Complete and efficient conversion of plant cell wall hemicellulose into high-value bioproducts by engineered yeast. Nat Commun 12: 4975

Trcek J, Mira NP, Jarboe LR (2015) Adaptation and tolerance of bacteria against acetic acid. Appl Microbiol Biotechnol 99: 6215–6229

Valgepea K, Adamberg K, Nahku R, Lahtvee PJ, Arike L, Vilu R (2010) Systems biology approach reveals that overflow metabolism of acetate in Escherichia coli is triggered by carbon catabolite repression of acetyl-CoA synthetase. BMC Syst Biol 4: 166

Warburg O, Wind F, Negelein E (1927) The metabolism of tumors in the body. J Gen Physiol 8: 519–530

Warnecke T, Gill RT (2005) Organic acid toxicity, tolerance, and production in Escherichia coli biorefining applications. Microb Cell Fact 4: 25

Wei N, Quarterman J, Kim SR, Cate JH, Jin YS (2013) Enhanced biofuel production through coupled acetic acid and xylose consumption by engineered yeast. Nat Commun 4: 2580

Weinert BT, Iesmantavicius V, Wagner SA, Scholz C, Gummesson B, Beli P, Nystrom T, Choudhary C (2013) Acetyl-phosphate is a critical determinant of lysine acetylation in E. coli. Molecular cell 51: 265–272

Wolfe AJ (2005) The acetate switch. Microbiol Mol Biol Rev 69: 12–50

Wolfe AJ, Chang DE, Walker JD, Seitz-Partridge JE, Vidaurri MD, Lange CF, Pruss BM, Henk MC, Larkin JC, Conway T (2003) Evidence that acetyl phosphate functions as a global signal during biofilm development. Mol Microbiol 48: 977–988

Xiao T, Khan A, Shen Y, Chen L, Rabinowitz JD (2022) Glucose feeds the tricarboxylic acid cycle via excreted ethanol in fermenting yeast. Nat Chem Biol

Yuan J, Doucette CD, Fowler WU, Feng XJ, Piazza M, Rabitz HA, Wingreen NS, Rabinowitz JD (2009) Metabolomics-driven quantitative analysis of ammonia assimilation in E. coli. Mol Syst Biol 5: 302

Zhang H, Pereira B, Li Z, Stephanopoulos G (2015) Engineering Escherichia coli coculture systems for the production of biochemical products. Proc Natl Acad Sci U S A 112: 8266–8271

