## Supplementary information for "From toxic waste to beneficial nutrient: acetate boosts *E. coli* growth at low glycolytic flux"

Contains Appendix Table S1, Figure S1, Figure S2.

| Primer name | Sequence 5′–3′ |
|---|---|
| **pgi forward external** | CTTCCAAAGTCACAATTCTCAAAATCAGAAGAGT |
| **pgi reverse external** | TGAACGCCTTATCCGGCCTAC |
| **pgi forward internal** | ACTTCGATGAAATGAAAGACGTTACGATCG |
| **pfkA forward external** | TTGTTATACTATTTGCACATTCGTTGGATCACT |
| **pfkA reverse external** | CATCGGTTTCAGGGTAAAGGAATCTG |
| **pfkA forward internal** | ATGATTAAGAAAATCGGTGTGTTGACAAGC |
| **acs forward external** | TACAGGTTTTGCGGGAGCAGCCGTTT |
| **acs reverse external** | CCTCATGCAGGACTTCATTATTAAGACGGTC |

**Appendix Table S1.** PCR primers used to check the *acs*, *pgi* and *pfkA* deletions.

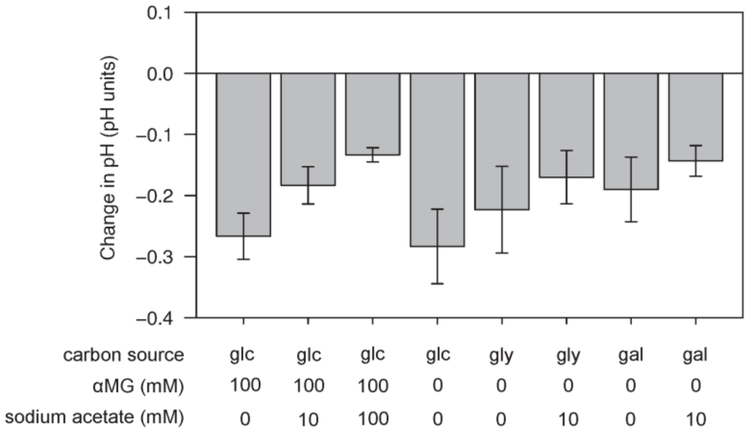


**Appendix Figure S1.** Change in pH of the cultivation medium during growth of *E. coli* K-12 BW25113 on glucose, glycerol or galactose (15 mM) plus different concentrations of αMG and acetate. Change in pH was measured in the exponential growth phase where growth rates and extracellular fluxes were measured (OD between ~0.1 and ~1.5). Mean values and standard deviations (error bars) were estimated from three independent biological replicates.

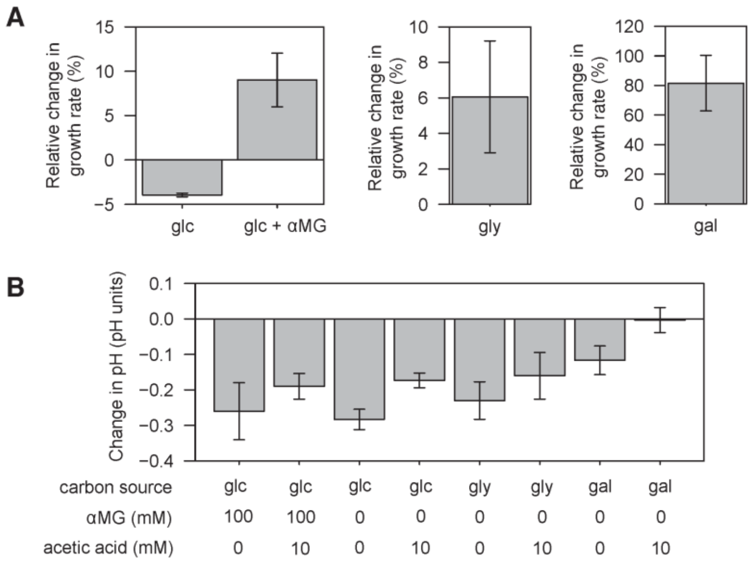


**Appendix Figure S2. Response of *E. coli* to acetic acid.**

A Relative change in the growth rate of *E. coli* K-12 BW25113 wild-type induced by the presence of 10 mM acetic acid during growth on glucose (without or with 100 mM αMG), glycerol or galactose. Mean values and standard deviations (error bars) were estimated from three independent biological replicates.

B Change in pH of the cultivation medium during growth of *E. coli* K-12 BW25113 wild-type on glucose, glycerol or galactose (15 mM) plus different concentrations of αMG and acetic acid. Change in pH was measured in the exponential growth phase where growth rates were measured (OD between ~0.1 and ~1.5). Mean values and standard deviations (error bars) were estimated from three independent biological replicates.
